# Vowel and formant representation in human auditory speech cortex

**DOI:** 10.1101/2022.09.13.507547

**Authors:** Yulia Oganian, Ilina Bhaya-Grossman, Keith Johnson, Edward F. Chang

**Affiliations:** Department of Neurological Surgery, University of California, San Francisco, 675 Nelson Rising Lane, San Francisco, CA 94158, USA; Center for Integrative Neuroscience, University Medical Center Tuebingen, Ottfried-Mueller-Str. 25, 72076 Tuebingen, Germany; University of California Berkeley, University of California, San Francisco Graduate Program in Bioengineering, Berkeley, CA 94720, USA; Department of Linguistics, University of California, Berkeley

## Abstract

Vowel sounds are a fundamental component of human speech across all languages. Vowels are cued acoustically by formants, the resonance frequencies determined by the shape of the vocal tract during speaking. An outstanding question in neurolinguistics is how the human brain processes vowel formants during speech perception. We used high-density intracranial recordings from the human speech cortex on the superior temporal gyrus (STG) while participants listened to natural continuous speech to address this question. We derived two-dimensional receptive fields based on the first and second formants to characterize tuning to vowel sounds. We found that neural activity at single STG sites was highly selective for particular zones in the formant space. Furthermore, this formant tuning shifted dynamically to adjust for speaker-specific spectral context. Despite this formant selectivity, local cortical responses were not sufficient to reliably discriminate between vowel categories. Instead, vowel category representations could be accurately decoded when using the entire population of formant encoding electrodes. Together, our results reveal that vowels are locally encoded in STG with complex acoustic tuning in two-dimensional formant space. As a population code this gives rise to phonological vowel perception.

## Introduction

Vowels are a significant component of all the world’s languages, and they play a critical role in our ability to comprehend spoken language (Fogerty & Humes, 2012; Fogerty & Kewley-Port, 2009). Vowel sounds are produced when vocal fold vibration is unobstructed, allowing a clear passage of air through the mouth, shaped by different positions of the jaw, tongue, and lips. For instance, the vowel sounds /i/ (as in “h<u>ee</u>d”) is produced with the tongue close to the front of the mouth, whereas /a/ (as in “h<u>a</u>d”) is produced with the tongue further back. These articulatory positions create acoustically distinct vowel sounds, defined by the value of the lowest two vocal tract resonance frequencies, which are known as the first and second formants (F1 and F2). Formants are considered the primary acoustic cues to vowel identity (Ladefoged & Johnson, 2014).

Sounds with different vowel identity can be very similar in absolute formant values when produced by different speakers. For example, /o/ (as in “h<u>oe</u>d”) produced by a speaker with a long vocal tract (and thus low voice) could have the same absolute formant values as /u/ (as in “wh<u>o’</u>d”) produced by a speaker with a short vocal tract (and thus high voice). As a result, the accurate mapping of formant values to vowel identity requires a normalization operation that computes formant frequencies relative to the speaker’s voice (Johnson & Sjerps, 2021). In sum, a rapid and correct mapping of formants to vowel identity relies on both the precise identification of the formant frequencies as well as their interpretation given the speaker context.

In linguistics, the mental representation of formants, specifically the independence of F1 and F2 and how this representation supports vowel normalization, has been long debated. One fundamental theory suggests that they are independently extracted and represented, mapping individual vowel sound segments onto coordinates in two-dimensional F1-F2 space (Nearey, 1989; Strange, 1989). To account for speaker context, proponents of this theory have advocated for an explicit normalization on both F1 and F2 values using features such as speaker pitch or higher formants. Alternatively, it has been suggested that vowel identification relies on the *relationship* between formants, such as the one-dimensional distance between F1 and F2 (Syrdal & Gopal, 1986). This theory of vowel perception is bolstered by the fact that the relationship between F1 and F2 is more consistent across speakers than the independent absolute values, thus implicitly accounting for speaker context (Miller, 1987; Peterson, 1961). Examining neural responses to vowel sounds in the human speech cortex provides us with a unique opportunity to dissociate these theoretical models and elucidate the mechanisms that underlie speaker-normalized vowel identification.

Several studies have shown that neural activity in the human auditory cortex is sensitive to vowel sounds and that it discriminates vowel identity in highly controlled acoustic contexts (Bonte et al., 2014; Formisano et al., 2008; Levy & Wilson, 2020). In the primary auditory cortex (PAC), a “tonotopic” area (Formisano et al., 2003) that is activated regardless of whether the presented stimulus is human speech (e.g. pure tones), this sensitivity is driven by narrow tuning to individual formant frequency bands (Hamilton et al., 2021; Khalighinejad et al., 2021). In contrast to PAC, in the human speech cortex on the lateral superior temporal gyrus (STG), the majority of neural representations are spectrally complex and broadband (Hamilton et al., 2021). Further, the STG shows stronger activity in response to speech than to other sound stimuli and this activity is more closely reflective of perceptual processing (Bhaya-Grossman & Chang, 2022; Yi et al., 2019). It thus remains an open question what neural representation in the STG underlies vowel perception in continuous, natural speech. Specifically, it is not yet clear whether formants are represented separately or in combination and further, whether this representation is tuned to narrow-band frequencies, possibly centered on single vowel categories (Shestakova et al., 2004), or broad formant ranges.

To address this, we utilized high-density direct intracranial recordings of neural activity from the surface of the human STG. The highly resolved spatial scale afforded by this recording technique was critical, as neighboring cortical sites that are just a few millimeters apart can differ significantly in their spectral tuning (Chang et al., 2010; Fox et al., 2020). The high temporal resolution of intracranial recordings allowed us to examine neural responses at the temporal scale of a single vowel sound. We used natural speech stimuli produced by a wide variety of speakers, which allowed us to record neural responses to a large set of vowel sounds, spanning the entire formant space.

With this approach we addressed four primary research questions. First, we asked how neural responses recorded at single electrodes in the STG were tuned to F1 and F2 in natural, continuous speech. We analyzed two-dimensional formant receptive fields of neural responses to determine whether neural tuning to F1 and F2 frequency ranges in human speech cortex is independent, and to characterize the mathematical properties of these tuning functions (Bertalmío et al., 2020; Fischer et al., 2009). Second, we asked how vowel category information can be extracted from the neural representation of formants using a population decoding approach. Third, we asked how and to what extent formant receptive fields in the STG are normalized for speaker properties. Finally, we used a controlled set of artificial vowel-like sounds extending beyond the natural vowel formant space with experimentally decorrelated F1 and F2 to definitively test the independence of formant encodings.

We found that neural responses on most single electrode sites in the STG were jointly tuned to both F1 and F2, resulting in heightened sensitivity to a specific zone within the vowel formant space. This sensitivity was non-linear and sigmoidal along each of the separate formant dimensions (F1 and F2). However, the location of heightened sensitivity did not coincide with single vowel category boundaries, and we did not find single electrode sites with selectivity for a single vowel category. Rather, single vowel categories could only be decoded at the population level, when information from differently tuned electrode sites was pooled together. Comparisons between neural responses produced by speakers with different vocal tract lengths showed that electrodes in the STG contain normalized, not absolute, formant representations, distinguishing it from the narrow-band frequency tuning in PAC. Though formant tuning on many active electrodes showed inverse tuning to the two formants (e.g. tuned to high F1 and low F2) when presented with natural speech, decorrelating the formants using a set of artificial vowels revealed a set of electrodes with the same direction of tuning to both formants (e.g. high F1 and high F2). This confirms that there exists a range of formant encoding types in the STG, and that F1 and F2 are neurally represented as coordinates in a two-dimensional formant space rather than as a ratio or distance.

## Results

### Cortical activity at single electrodes over human STG is sensitive to vowel differences

Nine Spanish monolingual patients (4 left hemisphere, see T1 for details), who were undergoing clinical monitoring for intractable epilepsy, volunteered to participate in this study. The Spanish vowel system is well suited to study vowel representation for multiple reasons: Spanish has only 5 vowel sounds (as opposed to e.g. up to 20 in some English dialects, (Hagiwara, 2004), that span a large range of formant values. Thus Spanish vowel categories are more easily separated in the acoustic formant space. As shown in Figure 1B and C, the median formant values for single vowel instances correspond strongly with vowel category (Fig. 1C category clustering: median silhouette score = 0.099, permutation test with 500 repetitions; p<0.002).

**Figure 1.**
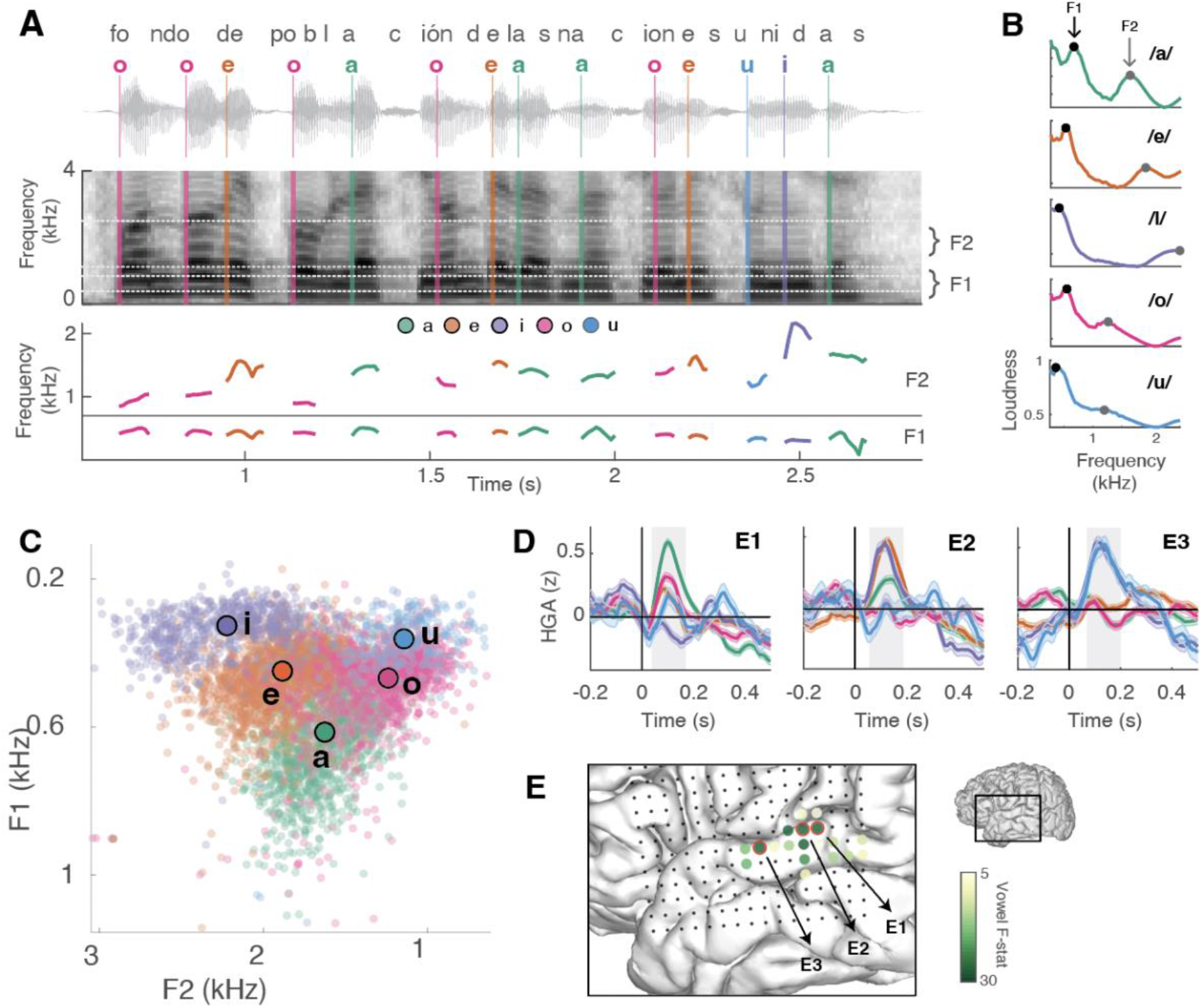
Neural activity on single electrodes in bilateral human STG is sensitive to vowels. **A.** Example Spanish stimulus sentence waveform (top), spectrogram (middle) and extracted formant trajectories (bottom). Vertical lines mark vowel onsets, colored by vowel identity. **B.** Median frequency spectrum for a single speaker per vowel category. Formant spectral peaks are marked in black and gray circles (F1 and F2 respectively). **C**. Median F1 and F2 values across all vowel instances in our stimulus set, colored by vowel identity. **D.** Differential average HGA responses to single vowel categories on three example STG electrodes. Error bars indicate standard error of the mean, gray shaded area marks time window of averaging for electrode selection. **E**. Vowel discriminative electrodes for a single participant. Electrodes cluster in the middle STG. Gray dots demarcate all grid electrodes, example electrodes from panel D are marked in red.

Moreover, while F1 and F2 tend to be inversely correlated across languages, this is less the case for Spanish than for English (Spanish r = −0.01, p = 0.31 in our stimulus set; English r = −0.21, p<0.0001 calculated from the stimulus set used in prior studies e.g. Mesgarani et al.,2014). While participants passively listened to 500 naturally produced Spanish sentences (Fig. 1A, vowels in the DIMEx corpus), we recorded neural activity from the lateral surface of the STG using high-density ECoG electrode grids. We extracted the analytic amplitude of neural responses in the high gamma range (HGA; 70 to 150 Hz), which is closely related to local neuronal firing and tracks neural activity at the temporal scales of natural speech (Mukamel & Fried, 2012; Ray & Maunsell, 2011).

We found that evoked HGA responses on a subset of STG electrodes discriminated between vowel categories (n = 125 of 291 speech responsive, range: 2-26 per participant, one-way peak F-statistic across vowel categories > 5) (see Fig. 1E for electrode distribution on the brain of an example participant). Responses peaked at about 100-150 ms post vowel onset, with different response magnitudes corresponding to different vowels. For example, in the prototypical electrode E1 (Fig. 1D top), responses were strongest for /a/ and weakest for /i/ sounds while in electrode E2 (Fig. 1D middle), responses were strongest for /e/ and /i/ and weakest for /u/ sounds. Finally in electrode E3, responses were strongest for /u/ and /i/ sounds and weakest for /e/ and /a/ sounds (Fig. 1D bottom).

Notably, none of these electrode responses show a preference for a single vowel category. That is, we did not observe instances where there was selectivity to a single vowel and no response to all other vowel sounds. In subsequent analyses, we include electrode responses that meet a vowel discriminability threshold (colored green in Fig. S1, maximal F-stat across vowel categories > 5, p<0.0001). Next, we characterized the formant representation on these electrodes in further detail.

### Non-linear monotonic tuning to vowel formant frequencies in human STG

We first asked how vowel formants are represented by vowel-discriminating populations. Specifically, we evaluated three alternative hypotheses: Narrow-band nonlinear frequency tuning centered on vowel categories (Fig. 2A left), linear monotonic encoding of an entire vowel formant range (Fig. 2A, middle panel), or nonlinear monotonic encoding of formants within a limited dynamic range (Fig. 2A, right panel). In the case of narrow-band frequency tuning, we expect neural populations to preferentially respond to a narrow range of formant frequencies, as is typical for frequency-tuning in primary auditory cortices (Bitterman et al.,2008; Hamilton et al., 2021). In particular, this model implies that the maximal neural response could be located in the center of the vowel’s formant range. In contrast, in monotonic formant encoding, we expect neural responses to increase across the range of possible formant values, with the maximal response located at the edges of the formant range. While linear encoding implies equal sensitivity to formant differences across the entire range, nonlinear encoding would result in heightened sensitivity to a narrower range of formant frequencies, and little to no sensitivity to frequencies outside of this range.

**Figure 2.**
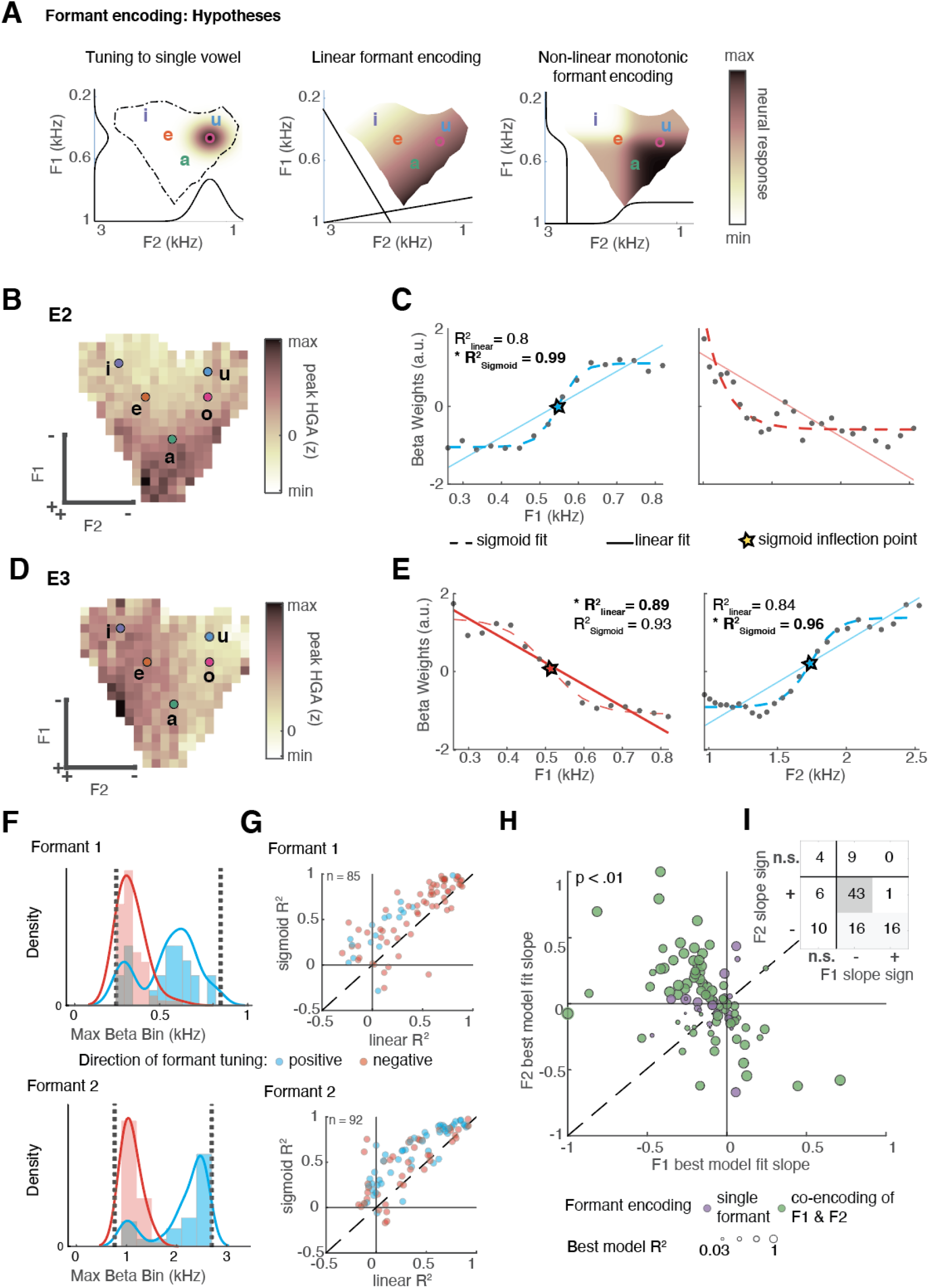
Non-linear monotonic encoding of vowel formant frequencies in human STG. **A.** Three alternative hypotheses for encoding of vowel formants on single electrodes. **B, D.** Example electrodes’ formant receptive fields. **C, E**. Formant tuning curves of the electrodes in B and D (F1 left, F2 right column). **F.** Frequency of maximal beta weights in F1 and F2 ranges shows that electrodes respond maximally at edges of formant range, as expected with monotonic encoding. **G**. Comparison between linear and sigmoid model fits for F1 (top) and F2 (bottom) shows sigmoidal encoding on a majority of sites. **H-I**. Distribution of slopes for F1 and F2 across vowel-responsive electrodes shows that most electrodes encode both F1 and F2, with a majority of the electrodes inversely encoding F1 and F2.

We tested which of these models captured neural activity best separately for every single electrode. First, to distinguish between narrow non-monotonic formant tuning and monotonic encoding, we examined whether neural responses peaked at the edges or in the center of the formant frequency range (Fig. 2F). Then, to discriminate between linear and non-linear encoding, we compared linear and sigmoidal models of neural responses using cross-validated R^2^, a measure of the portion of variance explained by each model. To estimate the neural response to vowel formants we used time-delayed ridge regression, also known as feature temporal receptive field modeling, F-TRF (Gill et al., 2006; Theunissen et al., 2001, Fig. S1). Model features of interest were the spectro-temporal content of the speech signal in the formant frequency ranges (F1: 250 - 800 Hz, F2: 1000 - 2500 Hz). The model produced a regression weight time series (beta weights) for each formant frequency bin. As confounding variables, we included sentence and vowel onsets, rising envelope edges, i.e. peakRate (Oganian & Chang, 2019), and speaker pitch predictors (Hamilton et al., 2021; Tang et al.,2017). For the main analysis, we identified the peaks in beta weight time courses (typically at 150ms across electrodes post vowel onset) and extracted mean beta weights in a 50ms window around it (125 - 175 ms). We then used these beta values to test the predictions of the three formant encoding models (frequency tuning, linear monotonic, non-linear monotonic) and to test whether each electrode showed tuning to only one or both formants.

For two representative electrodes E2 (Fig. 2C) and E3 (Fig. 2D), we found that responses were strongest towards one end of the vowel formant space, suggesting monotonic encoding of formant frequencies. On E2, response magnitudes were maximal for vowel tokens with low F1 and high F2 values (Fig. 2B): model beta weights increased with increases in F1 (Fig. 2C left), but decreased with increases in F2 (Fig. 2C right). Notably, for this electrode, F1 encoding was best described by a linear model (linear R^2^=0.89), whereas for F2 a non-linear, sigmoidal model fit its responses significantly better than a linear model (sigmoidal R^2^=0.96, linear R^2^=0.84). For example, in electrode E3, neural responses peaked for high F1 and low F2 values, with sigmoidal stimulus-response mappings in both formants (Fig. 2D,E).

Across all vowel-discriminating electrodes, we found that neural responses peaked near the boundaries of the vowel formant space, reflected in the bimodal distribution of maximal beta values for both formants (Fig 2F, F1: t(96) = 8.55, p<0.0001; F2: t(96) = 12.33, p<0.0001). We thus focused our attention on the comparison between linear and sigmoidal encoding of vowel formants. Across electrodes, we found that cross-validated R^2^ values were higher for the sigmoidal model than the linear model on 82.4% of electrodes for F1 and 81.5% of electrodes for F2. We found a mixture of preference for high and low formant values in both F1 and F2. This shows that STG neural populations have limited dynamic ranges: Each local population represents a subspace of the vowel formant space, such that a representation of the entire range emerges across the entire population.

Finally, we wanted to characterize the extent to which both formants are jointly encoded at a single electrode. Across electrodes, we found an inverse correlation between tuning to F1 and F2 (r = −0.448, p <0.0001). That is, electrodes with preference for high F1 values also prefered low F2 values, and vice versa, as previously found for an English language dataset (Mesgarani et al., 2014). Here, we found that this trend was driven by two main patterns. First, a majority of electrodes jointly encoded both formants (67 out of 80 electrodes), with a preference for negative tuning in F1 and positive tuning in F2 (41 out of 80 electrodes). In contrast, while a small subset of electrodes (n=10) had negative tuning to both formants, only a single electrode in our dataset had positive tuning to both formants. This raises the question: Does the lack of electrode sites with positive tuning to both formants reflect a specialization of STG for speech sound harmonics? Alternatively, it might be a confound of the limited formant frequency ranges in natural speech. We will address this question with Experiment 2 below.

Overall, these results show that neural activity at single electrode sites is only discriminative in a subspace of the overall vowel formant space. However, across electrodes, the range of formant tuning at the population level should be sufficient to represent the entire vowel space with a high fidelity. We directly test this in the next analysis step using population decoding.

### Discrimination between vowel categories emerges at the population level

We evaluated whether single vowel categories are represented at the population level by comparing the accuracy of vowel category decoding using different electrode subsets. All decoding was run using pairwise five-fold cross-validated support vector machine (SVM) classification on HGA from a time window of 50 ms around the median peak discriminability time point after vowel onset (see Methods for details on window selection).

First, we compared classifier accuracies derived from single electrodes versus from the entire electrode population (Fig. 3A). We found that decoding from the entire electrode population was significantly more accurate when taking into account all pairwise comparisons (average improvement: 2.2 - 9.1% correct averaged across pairwise comparisons, t-test showed significant difference between all single electrodes versus the entire population accuracies with p<0.0001, Fig. 3A). Notably, the best single electrodes did not necessarily show selective responses to a single vowel category (Fig. 3A right panel).

**Figure 3:**
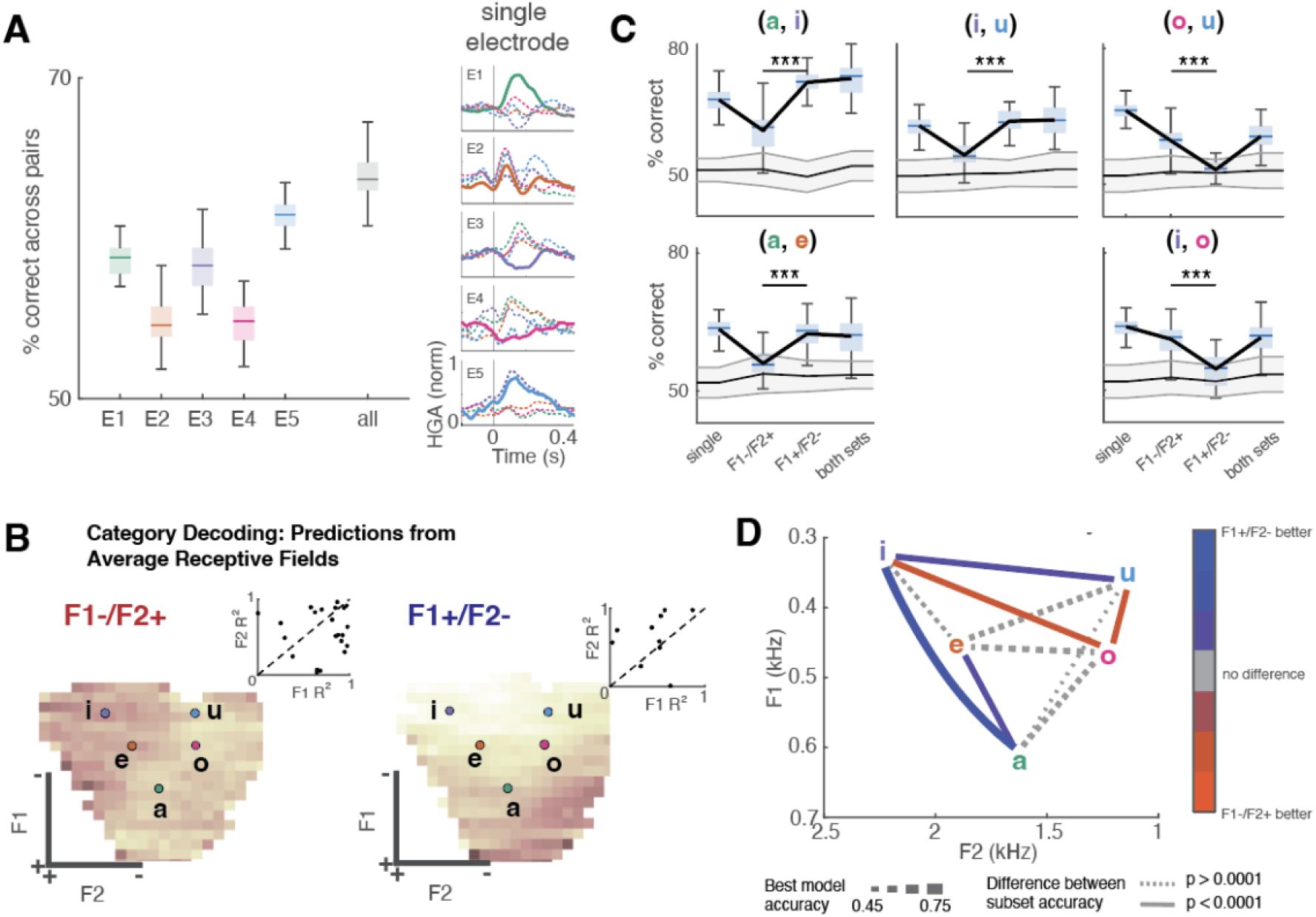
Emergent vowel representation at the population level. **A.** Vowel decoding accuracy from the best single electrode per vowel category (five unique electrodes across four subjects) as compared to the decoding accuracy using the electrode population. All accuracies averaged across ten pairwise comparisons. Right is the average HG response to vowel categories in five electrodes used in single electrode category decoding in the leftmost panel. Solid line corresponds to the vowel category for which the selected electrode shows best average accuracy, dashed lines correspond to all other categories. **B.** Average formant receptive fields for electrodes with different tuning types and corresponding F1/F2 sigmoidal model R^2^. **C.** Decoding accuracy from five vowel pairs separated by electrode sub-group (best single electrode, each tuning type, both tuning types; all with 50 repeats of resampled trials). Gray shaded bar indicates chance performance over repetitions. **D.** Summary of decoding accuracy across vowel pairs (all pairwise statistics in Table S6), color indicates the difference between decoding accuracy of the two electrode subsets (solid line marks significant difference). Line thickness corresponds to the magnitude of best model accuracy.

The improvement in classification accuracy from single electrode to the population could be due to an increase in the signal-to-noise ratio. However, this improvement may also reflect complementary vowel encoding. That is, single electrodes represent only select regions of the vowel formant range in high detail, and as a result, pooling information across electrodes with sensitivity to different select regions may be key to decoding vowel categories across all pairwise comparisons. To determine whether this was the case, we split electrodes into the two dominant tuning subsets, namely (F1-/F2+) and (F1+/F2-) type-electrodes. Formant receptive fields averaged across electrodes with the same directionality of F1/F2 tuning (F1− /F2+: n = 36 electrodes; F1+/F2-: n = 15 electrodes, Fig. 3C) suggested that each set would be critical for a subset of comparisons. For instance, average receptive fields suggested that F1+/F2-populations would be able to discriminate between /a/, /e/, and /i/, whereas this would not be the case for the F1-/F2+ population.

Decoding performed separately on the two subsets of electrodes confirmed this prediction, as can be seen in Fig. 3D for five exemplary comparisons (p<0.001 for significant comparisons between decoding accuracies on different subsets, see SOM T2 for details of pairwise comparisons). Moreover, this analysis clearly demonstrates that increased decoding accuracies at the population level reflect the addition of informational content rather than increases in signal-to-noise ratio. This is because accuracy of decoders based on the best electrode subset and both electrode subsets together did not significantly differ. Finally, Fig. 3D summarizes the accuracies of all pairwise comparisons, showing that the two subsets of electrodes discriminate different sets of vowel pairs. The F1-/F2+ electrodes show significantly higher accuracy for the /i/ - /o/ and /o/ - /u/ pairwise classification, while the F1+/F2-electrodes show higher accuracy for the /i/ - /a/, /i/ - /u/, and /e/ - /a/ classification. However, in conjunction, these two electrode sets contain the complementary information necessary to significantly discriminate between all vowel pairs.

### Shifting dynamic ranges underlie normalization of vowel representation for speaker vocal tract length

It is well established that the mapping between vowel formants and vowel categories depends on a speaker’s vocal tract length which is correlated with voice pitch (Johnson,2020; Johnson & Sjerps, 2021). Recent intracranial recordings have shown that neural responses in the STG to experimental vowel tokens adapt to speaker context, suggesting that the STG normalizes for speaker formant differences (Mesgarani et al., 2014; Sjerps et al., 2019). However, these studies did not explore the full range of vowels and formant frequencies in natural speech and therefore have not characterized the nature of neural tuning normalization in the STG. We thus wanted to determine whether the formant tuning of neural populations in STG shifts with speaker voice quality during listening to continuous speech. In line with prior work, we found that vowel formant frequencies increase with speaker pitch (Fig. 4A) and that normalizing for speaker pitch reduces the formant variance within vowel categories (Fig. 4B). We hypothesized that STG encoding of formants would reflect speaker-normalized rather than absolute formant frequencies. To test how speaker pitch affects the representation of vowel formants in human STG, we re-fit feature temporal receptive field models separately on subsets of data with low and high (>170 Hz) pitch levels, and estimate the sigmoidal fits to F1 and F2 model weights separately for each of the models. If STG representation of vowel formants reflects absolute formant frequencies, we expect to see no difference in formant tuning between models (Fig. 4C left). In contrast, if STG representations are normalized for speaker properties, we expect to see a shift in STG dynamic ranges, matching the shift in vowel formant space between single models (Fig. 4C right).

**Figure 4.**
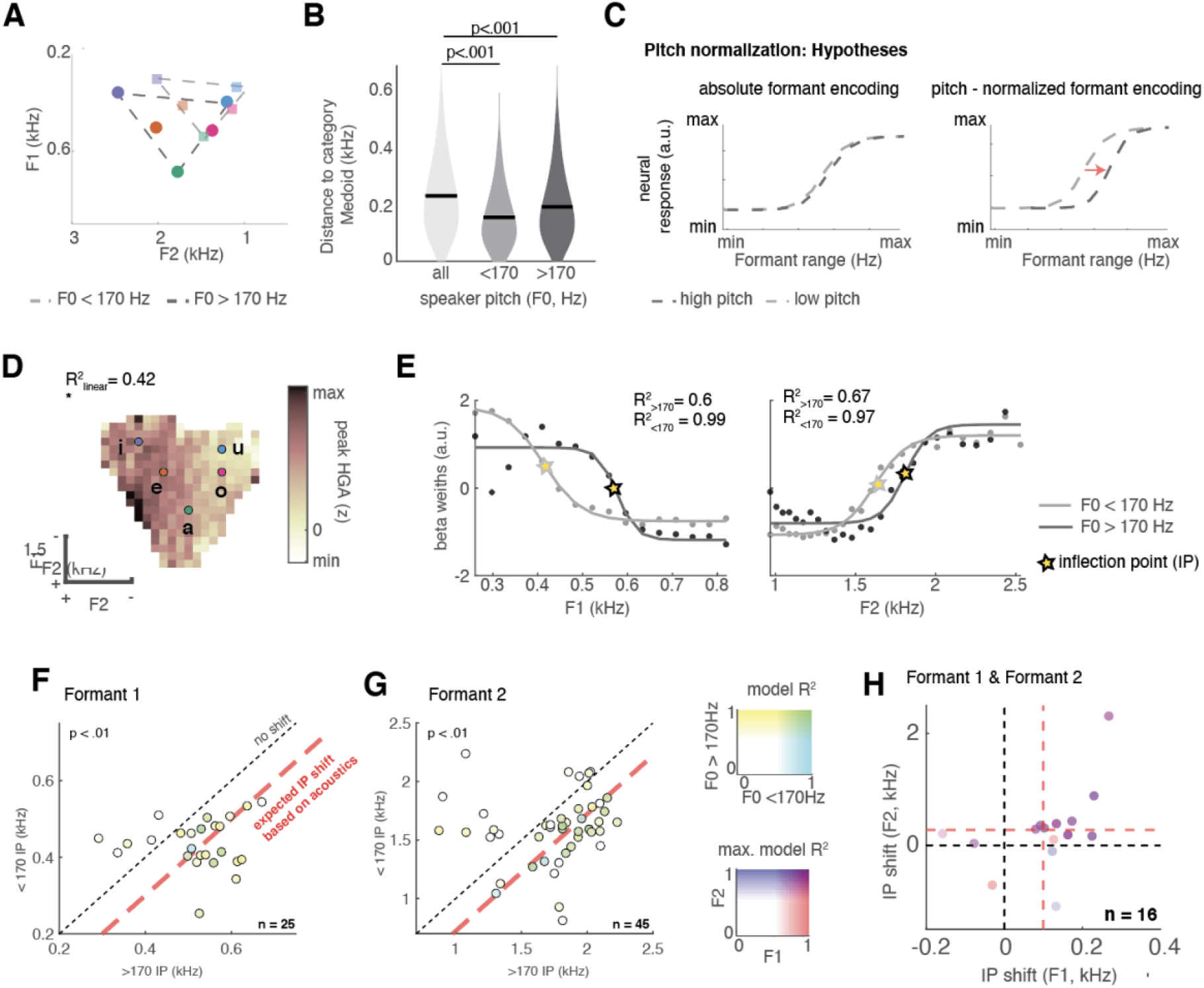
Shifting dynamic ranges underlie normalization of vowel representation for speaker pitch. **A.** Vowel formants shift between speakers with short and long vocal tracts (and thus high and low pitch). **B**. Formant normalization in two groups by high/low pitch reduces formant variability within vowel categories. **C.** Hypotheses for absolute (left) and pitch-normalized (right) encoding of vowel formants. **D.** Formant receptive field for an exemplary electrode. **E.** Formant frequency tuning for F1 (left) and F2 (right) on the same exemplary electrode, split by speaker pitch, shows a shift in tuning curves with increase in speaker pitch, as predicted by the normalization hypothesis. **F-G.** Tuning curve inflection points (IP), calculated separately for sentences with high and low speaker pitch, across all F1 (**F**) and F2 (**G**) encoding electrodes. Shift in IP below diagonal serves as an indicator for pitch normalization of formant tuning and is consistently present in electrodes with a good model fit for both pitch levels. **H**. IP shift for F1 and F2 on electrodes encoding both show that IP shifts on the majority of electrodes.

Fig. 4D-E shows tuning curves for low and high pitch speakers on a single example electrode. For this example electrode, neural responses shifted their dynamic ranges to higher frequencies in response to vowels produced by speakers with a high pitch, in line with speaker normalization for both F1 and F2. To quantify the magnitude and extent of this shift on STG across all electrodes with significant formant frequency tuning (n = 105), we focused on electrodes with robust frequency tuning curves for vowels produced by both high and low pitch speakers (F1 n = 25, F2 n = 45 (unique electrodes n = 68), 64.8% of electrodes). We extracted the sigmoidal fit inflection Points (IP) for each subset of speakers separately and assessed the difference in IP between models across electrode sites using linear mixed effects modeling (see methods for model details).

We found a systematic and robust shift of tuning curve inflection points on these electrodes towards higher formant ranges with increases in speaker pitch; F1 pitch t(111) = 3.288, SE = 10.426, p = 0.001; F2 pitch t(138) = 3.145, SE = 31.908, p = 0.002; see SOM for all fixed effects, Figure 4F, G). Remarkably, the average magnitude of the inflection point shift across electrodes mirrored the difference in average formant frequency between high and low pitch speakers (red lines in Fig. 4F, G). Finally, we also found that all electrodes with robust encoding of F1 and F2 also showed speaker normalization for both formants, Fig. 4H. Taken together, these analyses show that formant normalization for speaker voice characteristics is a general feature of formant encoding neural populations in the human STG.

### Vowel formant encoding emerges from general complex frequency tuning on STG

Our analysis of the representation of vowel formants in STG revealed that dynamic ranges on single electrode sites correspond to the range of formants found in natural speech. At the population level, we found that co-encoding of F1 and F2 with opposite direction of tuning was most frequent, as opposed to tuning for low or high values for both formants. This raised two questions. First, is this particular pattern of co-encoding, with opposite direction of preference for F1 and F2, due to the limited range and covariance between F1/F2 in natural speech? That is, when presented with complex harmonic sounds with a broader range of F1 and F2 values, will neural populations show sensitivity to formant values with the same direction of tuning? Second, does each formant value affect the neural responses independently (Fig. 5A top and middle row, joint independent encoding), or are co-encoding neural populations additionally integrating across F1 and F2, i.e. do responses to each formant depend on the value of the other formant (Fig. 5A bottom, joint interactive encoding)? Notably, the qualitative patterns differ between independent and interactive co-encoding: the latter shows a u-shaped tuning along one of the diagonals. Finally, we wanted to know whether STG encoding of the formant structure in sounds would continue outside the boundaries of the human vowel formant space. Alternatively, STG may contain separate neural populations encoding sound harmonic structure in speech and non-speech frequency ranges.

**Figure 5.**
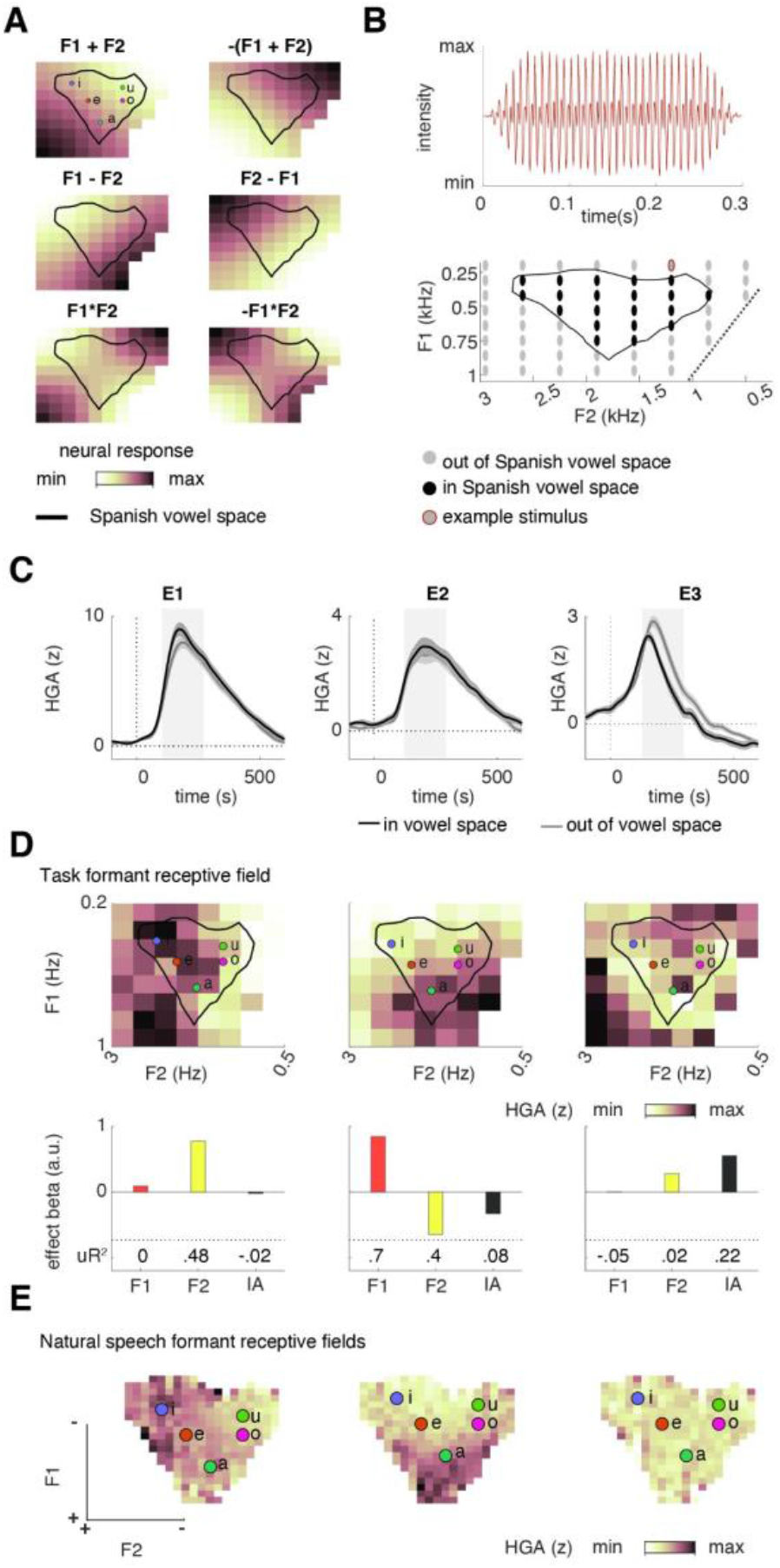
Vowel formant tuning emerges from complex frequency tuning on STG. **A.** Schematic of independent (top two rows) and interactive encoding of F1 and F2 (bottom row). **B.** Stimulus design for synthetic vowel task; Top: example token waveform. Bottom: F1 and F2 values task. Black dots mark tokens that fall within the natural vowel space, here defined by Spanish stimuli in our study. **C.** Responses to stimulus tokens that fall within and outside the Spanish vowel space on three example STG electrodes. Dark background marks peak area for averaging and further analyses. **D**. Formant receptive fields (top) and linear model effect R^2^ (bottom) for the same example electrodes. Black outline in formant receptive fields shows the vowel formant range in continuous Spanish sentences in the natural speech task. **E.** Formant receptive fields derived from DIMEx for the same example electrodes show similar response patterns as those derived from the synthetic vowel stimuli.

To test these questions, we designed a set of isolated artificial vowel-like sounds. Their F1 and F2 values covered the natural vowel space but also extended outside its range (10 values for F1, 10 values for F2, overall 86 unique tokens; Fig. 5B), with fixed pitch (F0 = 250 Hz) and higher formants (F3 = 3.1kHz, F4 = 3.3kHz, F5 = 4.135 kHz). Notably, this task was language-independent as vowels across languages fall within the same space due to physical constraints on vowel production. In Fig. 5B, the outline of the natural formant space is marked in black and derived from the Spanish stimuli used in Experiment 1. We chose to use isolated harmonic sounds rather than consonant-vowel combinations to reduce any effects of coarticulation and to ensure that tokens outside the vowel formant range could be perceived as non-linguistic. We recorded STG neural responses in a separate group of ECoG patients (n = 8, 2 Spanish monolingual, 6 English monolingual), while they passively listened to 9 - 20 repetitions of each synthetic vowel token. We found robust evoked HGA responses to single stimulus tokens on a subset of STG electrodes, which stereotypically peaked 150-200 ms after stimulus onset, in line with responses to natural speech. We first tested whether electrodes responded differently to tokens within and outside the natural formant range (black vs gray in Fig. 5B). Then, to model the encoding of F1 and F2, we extracted peak HGA for each stimulus token and evaluated the effects of F1, F2, and their linear interaction (F1*F2) on peak HGA magnitude. Across all 8 patients, analyses were focused on 160 electrodes (10-33 per patient), for which the best linear regression model explained at least 10% of total response variance.

Fig. 5C-E shows patterns of neural responses for three exemplary electrodes. Electrode E1 responded equally to tokens inside and outside the natural formant range (Fig. 5C left), with the strongest responses to high F2 values (Fig. 5D left), and no effect of F1 or the interaction term (Fig. 5D bottom). In contrast, responses on electrode E2 were highest for the combination of low F1 and high F2 values, as supported by the significant interaction term on this electrode (Fig. 5D middle). Finally, electrode E3 responded stronger to tokens outside the vowel formant range (Fig. 5C right), which was due to a significant positive interaction effect (Fig. 5D right). Notably, on all three electrodes the pattern of responses is generally smooth around the vowel space boundaries, suggesting that any difference between responses inside and outside this space are due to spectral and not speech tuning. Crucially, we found a high overlap between formant receptive fields derived from the synthetic vowel tokens and from natural speech stimuli, suggesting that both stimuli drive neural responses on STG to the same degree and in a comparable manner (Fig. 5E).

### Comparison between formant encoding in synthetic vowel sounds and in natural speech

Across all electrodes, we found that the main effects of F1 and F2 explained the most unique variance on single electrodes (F1: R^2^ median=0.07, max=0.71; F2: R^2^ median=0.07, max=0.72), with a minor but significant contribution of interaction terms (F1*F2: R^2^median=0.03, max=0.32; Fig. 6A). Notably, effect magnitudes for F1 and F2 were not correlated (r =-0.1, p=0.2 n.s. Fig. 6B), suggesting that each contributes independently to neural responses. In contrast, main and interaction effect magnitudes were negatively correlated (r=-0.52, p<0.001, Fig. 6C). That is, electrodes with large interaction effects had little independent contribution of F1 and F2 main effects and low R^2^ overall. This suggests that encoding of F1 and F2 and integration across formants are implemented by distinct STG populations with independent joint encoding of F1 and F2 dominating neural response patterns.

**Figure 6.**
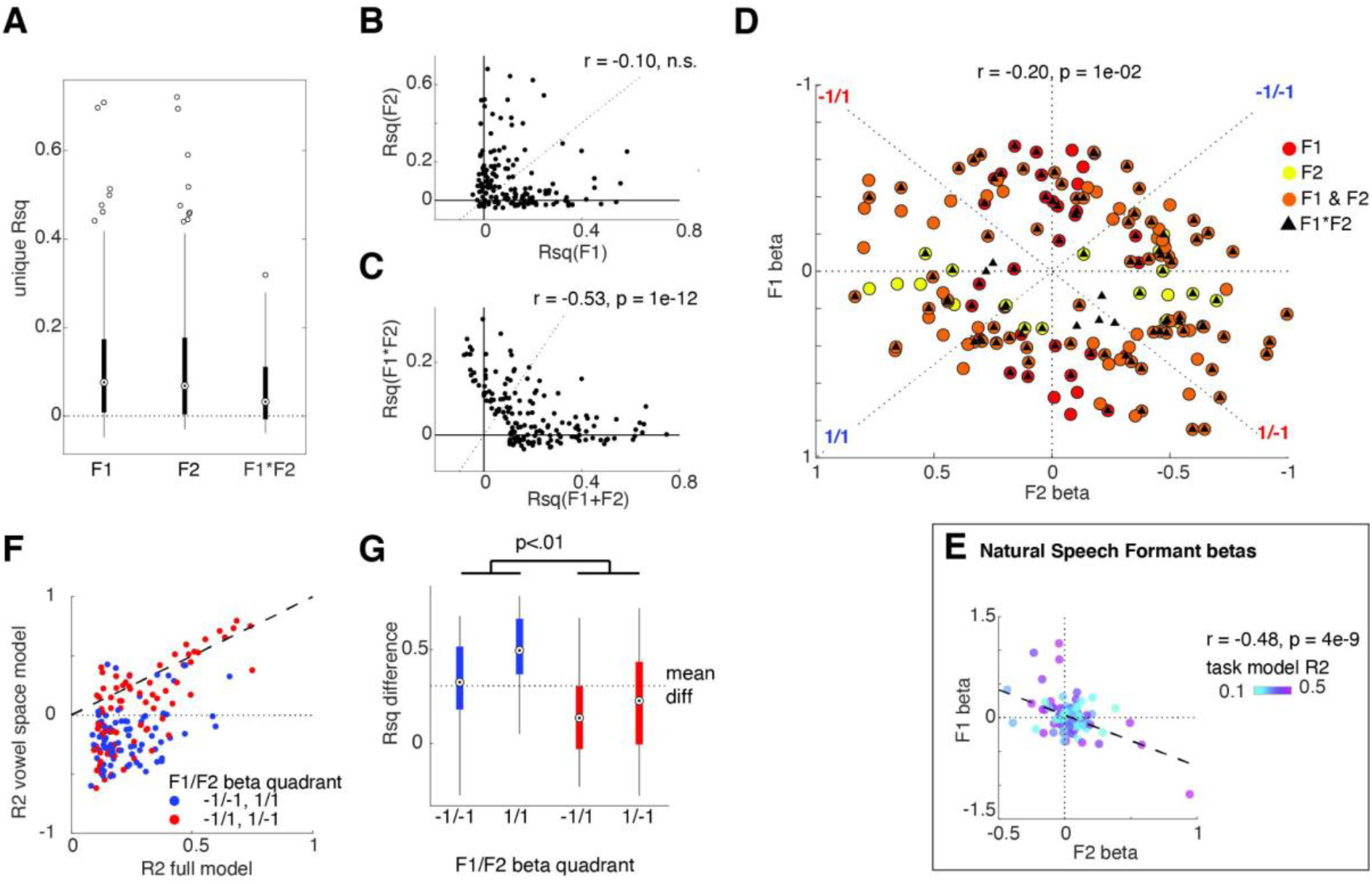
Comparison between formant encoding in synthetic vowel sounds and in natural speech. **A.** Effect R^2^ distribution across electrodes. **B-C.** Correlation between effect R^2^ shows that main effects of F1 and F2 are independent of each other (**B**), but are negatively correlated with the magnitude of the interaction effect (**C**). **D-E**. Across all electrodes, direction of effects for F1 and F2 is less correlated in the synthetic vowel task (**D**) than in the natural speech corpus (**E**). **F**. Model R^2^ for model fit on the entire vowel task stimulus set vs only on stimuli with formants located within the natural formant space. **G**. R^2^ values are overall lower in the reduced model, but this effect is stronger for electrodes with the same tuning directionality in F1 and F2.

In a second step, we asked whether STG representation of vowel formants is tailored to formant ranges found in natural speech. In particular, analyses of natural speech processing showed a negative correlation between the effects of F1 and F2 on neural responses to vowels in natural speech stimuli. In contrast, with our synthetic vowel stimuli we found only a weak negative correlation between F1 and F2 model weights (r = −0.2, p=0.01, Fig. 6D). Importantly, the range of tuning to different combinations of F1 and F2 values (N +/+, −/−, +/−, −/+ tuning direction) in our data speaks in favor of independent co-encoding of F1 and F2, rather than differential joint encoding.

We hypothesized that this discrepancy was due to the naturally limited range of F1 and F2 values in natural speech. To test this, we reran our analyses on a subset of stimuli with formants falling within the natural formant frequency range (as marked in Fig. 5B). Figure 6F shows the model R^2^ for full and subset models by electrode formant tuning. We found that in addition to the expected overall lower model R^2^ with a subset of stimuli, this reduction was more pronounced for electrodes with the same direction of tuning for both formants (Fig. 6G, p<0.01). Overall, this shows that populations with the same directionality of tuning for both formants are not as strongly activated by vowels but rather tuned to other harmonic sounds.

## Discussion

Here, we provide a comprehensive account of the representation of vowels in the human speech cortex on lateral superior temporal gyrus (STG). We found that local electrode sites in the STG encode vowels in complex two-dimensional receptive fields, defined jointly by the first two formant frequencies, and normalized for speaker voice characteristics. In each formant range, single electrode sites showed nonlinear, monotonic frequency tuning, with high sensitivity to a specific zone within the natural formant space. Because the spectral location of this zone did not correspond to any single vowel category, discrimination between vowel categories at single electrode sites was not reliable. However, vowel categories could be decoded from the neural response pattern across electrode sites. Furthermore, the analysis of neural responses to artificial vowel-like sounds with de-correlated F1 and F2 values showed that the complex tuning to formants in the STG independently represents F1 and F2, and extends beyond the natural vowel formant range.

Based on early work in phonetics, prior studies of cortical vowel encoding speculated that neural populations preferably encode the distance between F1 and F2 (F1-F2) (Syrdal & Gopal, 1986). While the neural responses to natural speech in this paper and earlier ECoG studies (Mesgarani et al., 2014) seemingly support this hypothesis, results from our synthesized vowel experiment suggest that this interpretation may have been oversimplified given the limited range of formants in natural speech. Using formant combinations that extend beyond the range that exist in natural speech, we show that co-encoding of F1 and F2 on individual electrodes in the middle STG spans all possible combinations of formant tuning, including sites with sensitivity for only one of the two formants. However, we also find that responses along the F1-F2 continuum contribute most to the discrimination of vowels, which reflects the main source of variance in the acoustic vowel space.

We also find that STG represents the spectral structure of vowels relative to the speaker rather than in absolute frequency terms (Johnson, 2020; Sjerps et al., 2019). While speaker normalization may reflect a general, speech-independent mechanism, a pure spectrotemporal model is incapable of making such a prediction. Context normalization can rely on multiple different cues in the stimulus: Here we used natural speech stimuli, where several cues to speaker normalization naturally co-occur (e.g. Johnson, 1988). While we rely on speaker pitch for our models, other co-varying cues are likely also involved, such as the distribution of the first two formant values for a given speaker (Sjerps et al., 2019) as well as the ratio between third formant and the first two (Monahan & Idsardi, 2010). Crucially, our results contradict recent work that argued that neural representations in the lateral STG reflect the veridical spectrotemporal content of sounds (Daube et al., 2019). This discrepancy is likely at least partly due to the different spatial resolution in our (intracranial recordings) and that work (scalp recordings). Overall, our work shows that STG representations of vowel formants are context-sensitive, leaving it for future work to pinpoint the neural sources of contextual information.

Together, these results directly address the linguistic debate regarding vowel representation (Strange, 1989). We find that the human speech cortex independently represents both vowel formants, rather than their relative distance (Syrdal & Gopal, 1986). This representation is normalized for speaker pitch (Fujisaki & Kawashima, 1968) with the magnitude of normalization matching the magnitude of the distance between low and high pitched speakers in the formant space. Speaker normalization may thus occur independently for each formant, based on separately calculated normalization cues.

It is well known that tonotopically organized neural populations in subcortical and primary auditory cortical areas represent the spectral structure of sounds and that, as a result, formant frequencies can be decoded from neural activity in these areas (Fisher et al., 2018;Obleser et al., 2006; Ohl & Scheich, 1997). In contrast, narrow frequency tuning and tonotopy have not been documented in the lateral STG (Hamilton et al., 2021), leaving it unclear how vowel sounds are represented in this area. Here, we show that vowel-encoding neural populations in the STG jointly encode both vowel formants, with nonlinear, sigmoidal tuning along each separate formant dimension. This tuning pattern allowed for heightened sensitivity to specific zones within the vowel formant space on single sites, which pooled into a representation of the entire space at the population level. Notably, our artificial vowel data showed that such representations also support the encoding of other harmonic sounds in the environment with formant ranges outside the vowel formant range. The specialization for different harmonic ranges may underlie recent findings of distinct neural populations for speech and music in this area (Boebinger et al., 2021; Norman-Haignere et al., 2015).

We found no evidence for focal representation of single vowels. That is, while tuning on local electrode responses represented parts of the two-dimensional formant space, it did not align with single vowel tunings. However, decoding of single vowels robustly emerged at the population level. This is in line with prior work focused on phonetic features in consonants, where results showed that local electrode responses on the STG encode acoustic-phonetic features, and that phonemes can only be extracted at the population level (Chang et al.,2010; Fox et al., 2020; Mesgarani et al., 2014). Our results build upon these findings, providing additional evidence that the holistic representation of speech sounds in the STG is organized as a distributed, spatial code. Specifically, we find that variation in the direction of formant co-encoding falls into two types of spatially interspersed electrode sites that each contribute to the discriminability of a subset of vowel categories. Such nonlinear encoding of vowel formants is a prerequisite and may be the basis for categorical vowel perception (Iverson & Kuhl, 1995; Kuhl, 1991; Levy & Wilson, 2020).

Our results demonstrate the broad and complex tuning of local electrode sites on the STG to the formants in natural vowel sounds. This tuning is best described by nonlinear, two-dimensional formant receptive fields that adapt to speaker characteristics. While a majority of local electrode sites jointly encode the first two formants, without a strong preference for a single vowel, vowel category labels can be extracted from the neural responses if they are aggregated across the population. Overall, we provide a comprehensive account of the representation of vowels in non-tonotopic areas of the auditory parabelt that are instrumental to the sensory processing of speech.

## Acknowledgements

We thank Will Schuermann, and Matthew K. Leonard for discussions of the manuscript. We thank Ben Speidel for help with anatomical localization of electrodes. We thank all members of the Chang Lab for help with the data analysis.

## Author Contributions

Conceptualization, YO, EFC; Methodology, YO, IBG, KJ; Software, YO, IBG; Formal Analysis, YO, IBG; Investigation, YO, IBG, KJ, EFC; Data Curation, YO, IBG; Writing - Original Draft, YO, IBG; Writing - Reviewing and Editing, YO, IBG, KJ, EFC; Visualization, YO, IBG, EFC; Supervision, YO, EFC; Project Administration, YO; Funding Acquisition, EFC.

## Methods

### Participants

15 (7 female) patients were implanted with 256-channel, 4-mm electrode distance, subdural ECoG grids as part of their treatment for intractable epilepsy. Electrode grids were placed over the peri-Sylvian region of one of the patients’ hemispheres (7 left hemisphere grids), as determined by clinical assessment. All participants had normal hearing and left-dominant language functions. The study was approved by the University of California, San Francisco Committee on Human Research. All participants gave informed written consent before experimental testing. Nine Spanish-native speakers (5 male, 4 LH) with little to no knowledge of English participated in the DIMEx corpus speech experiment.

Additionally, 8 participants (2 Spanish-native; 6 English-native, 7LH) listened to the synthesized vowel tokens of Experiment 2 (SOM T1). Participants in the synthesized vowel experiment also listened to the DIMEx corpus.

### Neural data acquisition and preprocessing

We recorded ECoG signals with a multichannel PZ2 amplifier, connected to an RZ2 digital signal acquisition system [TuckerDavis Technologies (TDT), Alachua, FL, USA], with a sampling rate of 3052 Hz. The audio stimulus was split from the output of the presentation computer and recorded in the TDT circuit time aligned with the ECoG signal. In addition, the audio stimulus was recorded with a microphone and also input to the RZ2. Data were online referenced in the amplifier. No further re-referencing was applied to the data.

Electrode positions were extracted from postimplantation computer tomography scans, coregistered to the patients’ structural magnetic resonance imaging and superimposed on three-dimensional reconstructions of the patients’ cortical surfaces using a custom-written imaging pipeline (Hamilton et al., 2017). Freesurfer was used to create a 3d model of the individual subjects’ pial surfaces, run automatic parcellation to get individual anatomical labels, and warp the individual subject surfaces into the cvs_avg35_inMNI152 average template.

Offline preprocessing of the data included (in this order) downsampling to 400 Hz, notch-filtering of line noise at 60, 120, and 180 Hz, exclusion of bad channels, and exclusion of bad time intervals. Bad channels were defined by visual inspection as channels with excessive noise. Bad time points were defined as time points with noise activity, which typically stemmed from movement artifacts, interictal spiking, or non-physiological noise. From the remaining electrodes and time points, the analytic amplitude in the high-gamma frequency range (70 to 150 Hz, HGA) was extracted using eight band-pass filters [Gaussian filters, logarithmically increasing center frequencies (70 to 150 Hz) with semi-logarithmically increasing bandwidths] with the Hilbert transform. The high-gamma amplitude was calculated as the first principal component of the signal in each electrode across all eight high-gamma bands, using principal components analysis. Last, the HGA was downsampled to 100 Hz and z-scored relative to the mean and SD of the data within each experimental block. All further analyses were based on the resulting time series.

### Experiment 1: Continuous speech (DIMEx)

#### Stimuli and procedure

Participants passively listened to a selection of 500 Spanish sentences from the DIMEx corpus (Pineda et al., 2004, 2010), spoken by a variety of native Mexican-Spanish speakers. Data in this task were recorded in five blocks of approximately 7-min duration each. Four blocks contained distinct sentences, and one block contained 10 repetitions of 10 sentences. Sentences were 2.5 to 8.03 s long and presented with an intertrial interval of 800 ms. The repeated block was used for validation of temporal receptive field models (TRF; see details below).

All stimuli were presented at a comfortable ambient loudness (~70 dB) through free-field speakers (Logitech) placed approximately 80 cm in front of the patients’ head using custom-written MATLAB R2016b (MathWorks, www.mathworks.com) scripts. Speech stimuli were sampled at 16000 Hz for presentation in the experiment. Participants were asked to listen to the stimuli attentively and were free to keep their eyes open or closed during the stimulus presentation.

Stimulus spectrograms were calculated using the NSL toolbox (http://nsl.isr.umd.edu/downloads.html) for Matlab. Continuous formant values were extracted using the praat software (Boersma & Weenink, n.d.),https://www.fon.hum.uva.nl/praat/,). We found that median formant values discriminate between vowel categories with a high accuracy. Thus all depictions of vowel tokens in two dimensional formant space reflect the token’s median formant values.

#### Analysis

##### Electrode selection

Analyses included electrodes located on the STG, for which the per electrode peak HGA response (over a contiguous window of 50 ms) after vowel onset significantly discriminated between vowel categories (using a one-way F-test and a non-corrected threshold of p < 0.001). Finally analyses included 122 electrodes, 2 to 26 within single patients (median = 12). For each of the below analysis, a subset of electrodes from this set were selected based on relevant criteria.

##### Feature temporal receptive field analysis (F-TRF)

We fit neural data with a linear temporal receptive field (F-TRF) model with different sets of speech features as predictors. In this model, the neural response at each time point [HGA(t)] is modeled as a weighted linear combination of features (f) of the acoustic stimulus (X) in a window of 600 ms before that time point, resulting in a set of model coefficients, b_1…, d_ (Fig. S1) for each feature f, with d = 60 for a sampling frequency of 100 Hz and inclusion of features from a 600 ms window (See previous work, Mesgarani et al., 2014).

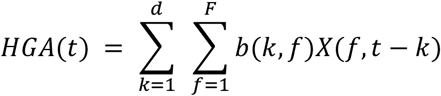

The models were estimated separately for each electrode, using five-fold cross-validation (80% train, 20% test). The regularization parameter was estimated using a 10-way bootstrap procedure on the training dataset for each electrode separately. Then, a final value was chosen as the average of optimal values across all electrodes for each patient. For all models, predictors and dependent variables were z-scored and scaled to between −1 and 1 before entering the model. This approach ensured that all estimated beta values were scale free and could be directly compared across predictors, with beta magnitude interpreted as an index for the contribution of a predictor to model performance.

Predictors in the model included spectral ranges that spanned the 5-95th percentile of frequency values for the first two formants for all vowels, as well as sentence onsets, vowel onsets, and predictors for timing and magnitude of peakRate, a marker of speech envelope rise time (for peakRate feature structure see Oganian & Chang, 2019). Vowel and sentence onset predictors were timed to onsets of the respective phonemes in the speech signal (see Fig. S1 for procedure diagram).

##### Linear and sigmoidal model comparisons on F-TRF betas

Linear, sigmoidal, and quadratic curves were fitted to the mean beta weights around the beta peak in a 600 ms window (125-175ms), as estimated by the F-TRF model described above. Curve fitting for all model types was leave-one-out cross-validated (16 and 26 times for F1 and F2 bins respectively). This allowed us to directly compare models with different numbers of free parameters. Corresponding model R^2^ values were calculated based on the average error derived from the cross-validated predictions. R^2^ values were calculated as 1 - RSS/SST, where RSS is defined as the residual sum of squares and SST is defined as the total sum of squares. Note, this coefficient of determination measure (R^2^) could be negative if the regression predictions are further from the true value than a model that predicts the sample average. Bayesian Information Criterion (BIC) was used to validate the R^2^ values. According to this metric, the sigmoidal model outperformed the linear model on 31.4 % of electrodes for F1 and 25.7% of electrodes for F2.

##### Vowel category decoding

Binary SVM linear classifiers (using pre-built function from Matlab Statistics and Machine Learning Toolboxes) were trained on neural data to predict pairwise vowel categories from a fixed window of 50 ms around the time point of peak discriminability between vowel categories (centered at approximately 150 ms post-vowel onset). Four types of classifiers were constructed: population level classifiers, two sets of population-subset classifiers, and single electrode classifiers. Population level classifiers were trained and tested on the output of principal component analysis (PCA) applied to neural activity from a population of electrodes (n = 54) spanning 4 subjects with sufficiently overlapping stimulus sets. To ensure that the pairwise category accuracy was comparable across pairs, data were subset to contain equal numbers of samples per vowel category, equal to the number of samples in the least frequent vowel category (/u/) resulting in approximately 110 samples per vowel category. Population-subset classifiers were also trained and tested on the output of a PCA and included electrodes with either F1+/F2-(n = 40) or F1-/F2+ (n = 14) individual tuning profiles derived from previous analysis. Single electrode classifiers were trained and tested on the neural activity recorded at single electrodes (n=105). Each classifier was 5-fold cross-validated and reported accuracy measures were averaged across each of the cross-validation sets. Significance testing was based on classification accuracies for data with permuted vowel category labels.

##### Speaker normalization

To determine the extent to which cortical responses to formants depend on speaker physiology, separate F-TRF models were were fit to neural data using subsets of speakers with an average fundamental frequency across the sentence of either less than or greater than 170 Hz, using the same predictors as the model described above. A total of 233 sentences (4840 vowel instances) were used in the low pitch speaker model and 267 sentences (5750 vowel instances) were used in the model corresponding to high pitch speakers. Curve fitting was performed on the speaker subset F-TRF model beta weights in the same way as described above. Inflection point shifts were calculated using the parameters derived from each of the F-TRF model types. Effects of speaker subset on inflection point shift was determined using linear mixed effects modeling (using the Matlab Statistics and Machine Learning Toolbox) with fixed effects of model R^2^ and speaker, and random intercepts and slopes for speaker within electrode (Inflection Point Position ~ Model R^2^ * Speaker + (1+ Speaker | Electrode)).

### Experiment 2: Synthetic vowels

#### Stimuli and procedure

Participants passively listened to a set of synthesized vowel tokens. Stimuli were synthesized using an online version of the Klatt vowel synthesizer (www.source-code.biz/klattSyn/), with fixed pitch (250 Hz), F3 (3.1 kHz) and F4 (3.3 kHz). F1 and F2 were varied orthogonally to cover the entire vowel formant range as well as values outside those occurring in natural speech. For F1, we selected 10 values between 200 and 100 Hz (200, 255, 310, 420, 530, 640, 750, 860, 915, 970); for F2 we selected 10 values between 500 and 3000 Hz (500, 650,850,1200,1550, 1900, 2250, 2600, 2800, 2950). Stimuli covered all F1/F2 combinations with F2 larger than F1. Data in this task were recorded in five blocks of approximately 4-min duration each, resulting in 8 to 10 repetitions per token in each patient. Tokens were 300 ms long and were presented with an average intertrial interval of 800 ms, randomly sampled from a uniform distribution between 700 and 900 ms.

All stimuli were presented at a comfortable ambient loudness (~70 dB) through free-field speakers (Logitech) placed approximately 80 cm in front of the patients’ head using custom-written MATLAB R2016b (MathWorks, www.mathworks.com) scripts and psychtoolbox (Brainard, 1997; Kleiner et al., 2007; Pelli, 1997). Stimuli were sampled at 16 kHz for presentation in the experiment. Participants were asked to listen to the stimuli attentively and were free to keep their eyes open or closed during the stimulus presentation.

#### Analysis

##### Electrode selection

Analyses included electrodes located on the STG, which showed robust evoked responses to vowel stimuli, defined as electrodes for which the best linear model, either with only main effects or with main effects and interaction terms, explained at least 10 % of the variance. Analyses contained 160 electrodes, 10 to 33 within single patients.

##### Analysis of variance

As neural responses had a stereotypical evoked response peaking between 100 and 400 ms after stimulus onset, we focused our analyses on the mean HGA in that time window. Single time point analyses of the entire HGA time course produced qualitatively the same results. We modeled the main effects of F1, F2 and their linear interaction (F1*F2) onto evoked HGA responses to synthetic vowel onset using linear models. To assess the unique variance for main effects, we compared a main effect model (HGA ~ F1 + F2) to models containing only one formant (e.g., HGA ~ F1). To assess the unique variance explained by the interaction term, a full model (HGA ~ F1 + F2 + F1*F2) was compared to the model containing main effects only (HGA ~ F1 + F2).

For comparability across electrodes, predictors and HGA were z-scored prior to model fitting. For comparison to natural speech data, S-TRF models were fitted to natural speech response data on the same electrodes, using the same procedures as described above. To assess the effect of the formant range onto electrode response properties, models were fitted twice: First using all stimulus tokens, and second using only the subset of stimuli with formant values within the DIMEX vowel formant range.

##### Data and code availability

All code will be available for download via github upon publication. Data will be available from the authors upon request.

## Supplementary Materials

**Table S1.**
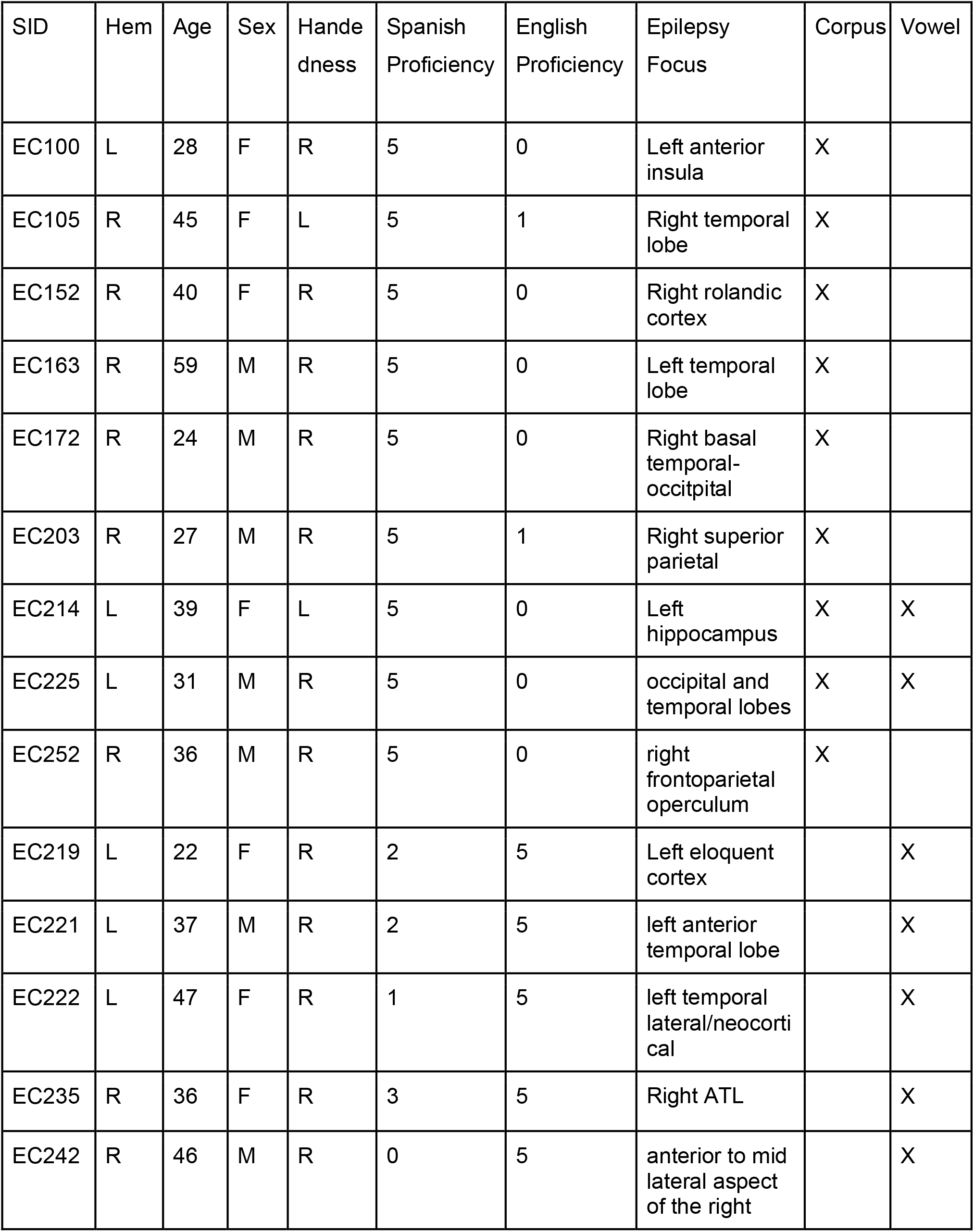
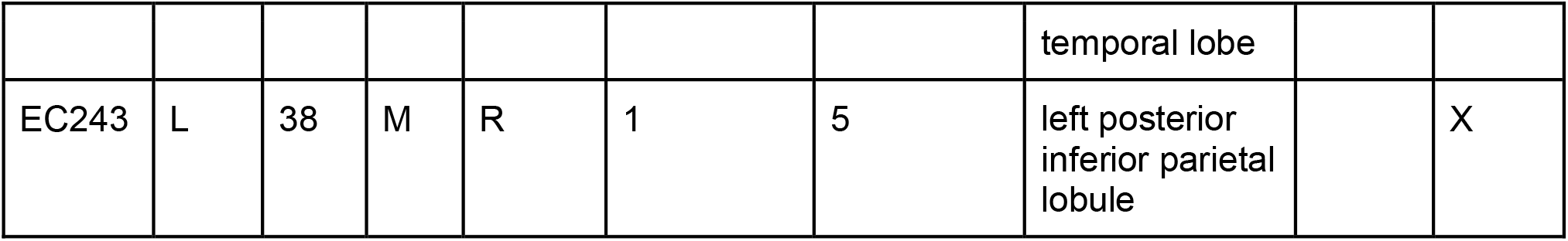
Participant details

| SID | Hem | Age | Sex | Handedness | Spanish Proficiency | English Proficiency | Epilepsy Focus | Corpus | Vowel |
| --- | --- | --- | --- | --- | --- | --- | --- | --- | --- |
| EC100 | L | 28 | F | R | 5 | 0 | Left anterior insula | X |  |
| EC105 | R | 45 | F | L | 5 | 1 | Right temporal lobe | X |  |
| EC152 | R | 40 | F | R | 5 | 0 | Right rolandic cortex | X |  |
| EC163 | R | 59 | M | R | 5 | 0 | Left temporal lobe | X |  |
| EC172 | R | 24 | M | R | 5 | 0 | Right basal temporal-occipital | X |  |
| EC203 | R | 27 | M | R | 5 | 1 | Right superior parietal | X |  |
| EC214 | L | 39 | F | L | 5 | 0 | Left hippocampus | X | X |
| EC225 | L | 31 | M | R | 5 | 0 | occipital and temporal lobes | X | X |
| EC252 | R | 36 | M | R | 5 | 0 | right frontoparietal operculum | X |  |
| EC219 | L | 22 | F | R | 2 | 5 | Left eloquent cortex |  | X |
| EC221 | L | 37 | M | R | 2 | 5 | left anterior temporal lobe |  | X |
| EC222 | L | 47 | F | R | 1 | 5 | left temporal lateral/neocortical |  | X |
| EC235 | R | 36 | F | R | 3 | 5 | Right ATL |  | X |
| EC242 | R | 46 | M | R | 0 | 5 | anterior to mid lateral aspect of the right temporal lobe |  | X |
| EC243 | L | 38 | M | R | 1 | 5 | left posterior<br>inferior parietal<br>lobule |  | X |

**Figure S1.**
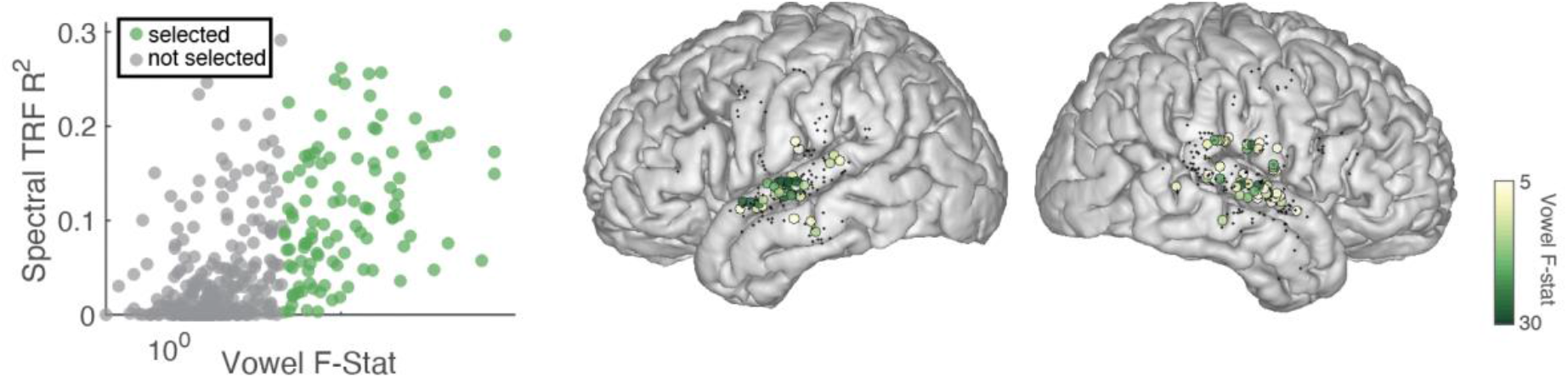
S1 TRF model vs. FSTAT to show selection criteria. Comparison of vowel discrimination metric (F-stat over peak-time period shown in leftmost panel) to an spectro-temporal receptive field (STRF) model R^2^, green electrodes are selected for subsequent analysis. Electrodes are pooled across nine Spanish monolingual participants (n = 125, 77.9% STG electrodes) and the rightmost panels show their anatomical location.

**Figure S2.**
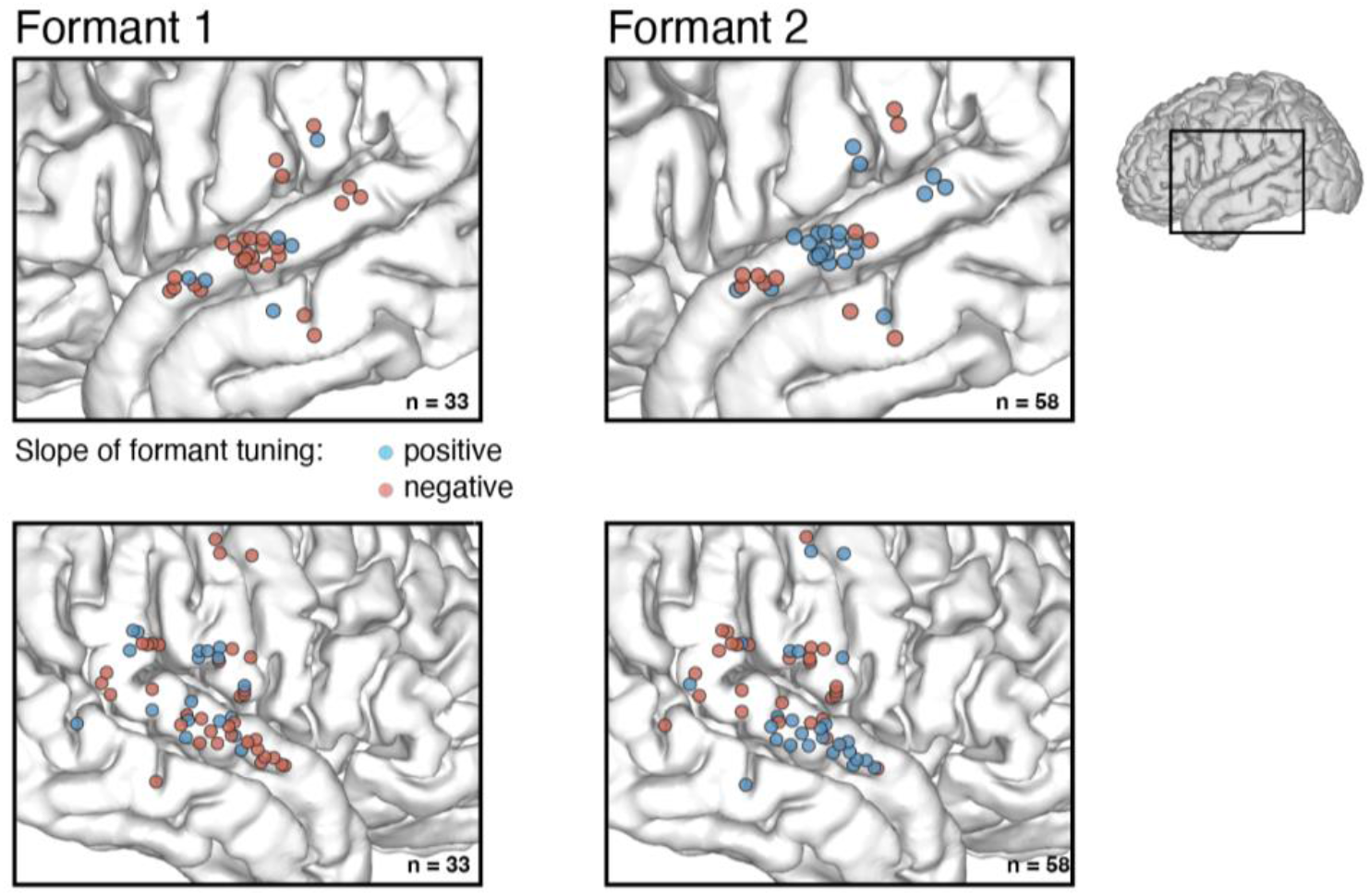
S2 F1/F2 Tuning on MNI Brain. Anatomical location of electrodes used in analysis for Fig. 2. Color indicates the directionality of formant tuning for either F1 or F2 across all 9 participants.

**Table S4.** Pitch effect on Inflection Point in F1. Infl ~ 1 + Pitch*Rsq + (1 + Pitch | Elec)

| predictor | estimate | SE | tstat | df | p | CI |
| --- | --- | --- | --- | --- | --- | --- |
| intercept | 496.04 | 11.19 | 44.4 | 111 | <0.001 | [473, 518] |
| pitch | 34.227 | 10.42 | 3.3 | 111 | 0.001 | [14, 55] |
| rsq | -11.79 | 10.12 | -1.2 | 111 | 0.24 | [-31, 8] |
| pitch*rsq | 3.04 | 10.0 | 0.3 | 111 | 0.76 | [-17, 23] |

**Table S5.** Pitch effect on Inflection Point in F2. Infl ~ 1 + Pitch*Rsq + (1 + Pitch | Elec)

| predictor | estimate | SE | tstat | df | p | CI |
| --- | --- | --- | --- | --- | --- | --- |
| intercept | 1711.1 | 35.9 | 47.6 | 138 | <0.001 | [1640, 1782] |
| pitch | 100.35 | 31.9 | 3.1 | 138 | 0.002 | [37, 163] |
| rsq | 48.02 | 33.9 | 1.4 | 138 | 0.16 | [-19, 115] |
| pitch*rsq | 51.66 | 32.8 | 1.6 | 138 | 0.12 | [-13, 116] |

**Table S6.** Pairwise Category Decoding Effects. (Two accuracies are median across 50 bootstrap reps for F1-/F2+ set and F1+/F2-set respectively)

| Pair | F1- Accuracy | F1+ Accuracy | Z-val | P-val |
| --- | --- | --- | --- | --- |
| a/e | 0.5596 | **0.6241** | -10.2591 | ** <0.0001 |
| a/i | 0.6172 | **0.7247** | -15.6310 | ** <0.0001 |
| a/o | 0.6092 | 0.6044 | 0.6575 | 0.5124 n.s. |
| a/u | 0.5568 | **0.5754** | -2.6375 | ** 0.0097 |
| e/i | **0.5782** | 0.5583 | 2.8676 | * 0.0051 |
| e/o | 0.6360 | **0.6554** | -3.4965 | ** <0.0001 |
| e/u | 0.6275 | 0.6373 | -1.4368 | 0.1540 n.s. |
| i/o | **0.6115** | 0.5463 | 9.3097 | ** <0.0001 |
| i/u | 0.5656 | **0.6403** | -12.5062 | ** <0.0001 |
| o/u | **0.5963** | 0.5321 | 11.0080 | ** <0.0001 |

## References

Bertalmío, M., Gomez-Villa, A., Martín, A., Vazquez-Corral, J., Kane, D., & Malo, J. (2020). Evidence for the intrinsically nonlinear nature of receptive fields in vision. Scientific Reports, 10(1), 16277.

Bhaya-Grossman, I., & Chang, E. F. (2022). Speech Computations of the Human Superior Temporal Gyrus. Annual Review of Psychology, 73, 79–102.

Bitterman, Y., Mukamel, R., Malach, R., Fried, I., & Nelken, I. (2008). Ultra-fine frequency tuning revealed in single neurons of human auditory cortex. Nature, 451(7175), 197–201.

Boebinger, D., Norman-Haignere, S. V., McDermott, J. H., & Kanwisher, N. (2021). Music-selective neural populations arise without musical training. Journal of Neurophysiology, 125(6), 2237–2263.

Boersma, P., & Weenink, D. (n.d.). Praat: doing phonetics by computer. [Computer Program]. Retrieved August 23, 2022, from https://www.fon.hum.uva.nl/praat/

Bonte, M., Hausfeld, L., Scharke, W., Valente, G., & Formisano, E. (2014). Task-dependent decoding of speaker and vowel identity from auditory cortical response patterns. The Journal of Neuroscience: The Official Journal of the Society for Neuroscience, 34(13), 4548–4557.

Brainard, D. H. (1997). The Psychophysics Toolbox. Spatial Vision, 10(4), 433–436.

Chang, E. F., Rieger, J. W., Johnson, K., Berger, M. S., Barbaro, N. M., & Knight, R. T. (2010). Categorical speech representation in human superior temporal gyrus. Nature Neuroscience, 13(11), 1428–1432.

Daube, C., Ince, R. A. A., & Gross, J. (2019). Simple Acoustic Features Can Explain Phoneme-Based Predictions of Cortical Responses to Speech. Current Biology: CB, 29(12), 1924–1937.e9.

Fischer, B. J., Anderson, C. H., & Peña, J. L. (2009). Multiplicative auditory spatial receptive fields created by a hierarchy of population codes. PloS One, 4(11), e8015.

Fisher, J. M., Dick, F. K., Levy, D. F., & Wilson, S. M. (2018). Neural representation of vowel formants in tonotopic auditory cortex. NeuroImage, 178, 574–582.

Fogerty, D., & Humes, L. E. (2012). The role of vowel and consonant fundamental frequency,envelope, and temporal fine structure cues to the intelligibility of words and sentences. The Journal of the Acoustical Society of America, 131(2), 1490–1501.

Fogerty, D., & Kewley-Port, D. (2009). Perceptual contributions of the consonant-vowel boundary to sentence intelligibility. The Journal of the Acoustical Society of America, 126(2), 847–857.

Formisano, E., De Martino, F., Bonte, M., & Goebel, R. (2008). “ Who” is saying” what”?Brain-based decoding of human voice and speech. Science, 322(5903), 970–973.

Formisano, E., Kim, D. S., Di Salle, F., van de Moortele, P. F., Ugurbil, K., & Goebel, R. (2003). Mirror-symmetric tonotopic maps in human primary auditory cortex. Neuron, 40(4), 859–869.

Fox, N. P., Leonard, M., Sjerps, M. J., & Chang, E. F. (2020). Transformation of a temporal speech cue to a spatial neural code in human auditory cortex. eLife, 9. https://doi.org/10.7554/eLife.53051

Fujisaki, H., & Kawashima, T. (1968). The roles of pitch and higher formants in the perception of vowels. IEEE Transactions on Audio and Electroacoustics, 16(1), 73–77.

Gill, P., Zhang, J., Woolley, S. M. N., Fremouw, T., & Theunissen, F. E. (2006). Sound representation methods for spectro-temporal receptive field estimation. Journal of Computational Neuroscience, 21(1), 5–20.

Hagiwara, R. E. (2004). An Acoustic Analysis of Vowel Variation in New World English(review). In Language (Vol. 80, Issue 4, pp. 903–903). https://doi.org/10.1353/lan.2004.0191

Hamilton, L. S., Chang, D. L., Lee, M. B., & Chang, E. F. (2017). Semi-automated Anatomical Labeling and Inter-subject Warping of High-Density Intracranial Recording Electrodes in Electrocorticography. Frontiers in Neuroinformatics, 11, 62.

Hamilton, L. S., Oganian, Y., Hall, J., & Chang, E. F. (2021). Parallel and distributed encoding of speech across human auditory cortex. Cell, 184(18), 4626–4639.e13.

Iverson, P., & Kuhl, P. K. (1995). Mapping the perceptual magnet effect for speech using signal detection theory and multidimensional scaling. The Journal of the Acoustical Society of America, 97(1), 553–562.

Johnson, K. (1988). Processes of speaker normalization in vowel perception (M. Beckman (ed.)) [The Ohio State University]. https://www.proquest.com/dissertations-theses/processes-speaker-normalization-vowel-perception/docview/303700096/se-2

Johnson, K. (2020). The ΔF method of vocal tract length normalization for vowels. Laboratory Phonology, 11(1). https://doi.org/10.5334/labphon.196

Johnson, K., & Sjerps, M. J. (2021). Speaker Normalization in Speech Perception. In The Handbook of Speech Perception (pp. 145–176). Wiley. https://doi.org/10.1002/9781119184096.ch6

Khalighinejad, B., Patel, P., Herrero, J. L., Bickel, S., Mehta, A. D., & Mesgarani, N. (2021). Functional characterization of human Heschl’s gyrus in response to natural speech. NeuroImage, 235, 118003.

Kleiner, M., Brainard, D., & Pelli, D. (2007). What’s new in Psychtoolbox-3? https://pure.mpg.de/rest/items/item_1790332/component/file_3136265/content

Kuhl, P. K. (1991). Human adults and human infants show a “perceptual magnet effect” for the prototypes of speech categories, monkeys do not. Perception & Psychophysics, 50(2), 93–107.

Ladefoged, P., & Johnson, K. (2014). A course in phonetics. Nelson Education.

Levy, D. F., & Wilson, S. M. (2020). Categorical Encoding of Vowels in Primary Auditory Cortex. Cerebral Cortex, 30(2), 618–627.

Mesgarani, N., Cheung, C., Johnson, K., & Chang, E. F. (2014). Phonetic feature encoding in human superior temporal gyrus. Science, 343(6174), 1006–1010.

Miller, J. D. (1987). Auditory-perceptual interpretation of the vowel. In The Journal of the Acoustical Society of America (Vol. 81, Issue S1, pp. S16–S16). https://doi.org/10.1121/1.2024119

Monahan, P. J., & Idsardi, W. J. (2010). Auditory sensitivity to formant ratios: Toward an account of vowel normalisation. Language and Cognitive Processes, 25(6), 808–839.

Mukamel, R., & Fried, I. (2012). Human intracranial recordings and cognitive neuroscience. Annual Review of Psychology, 63, 511–537.

Nearey, T. M. (1989). Static, dynamic, and relational properties in vowel perception. The Journal of the Acoustical Society of America, 85(5), 2088–2113.

Norman-Haignere, S., Kanwisher, N. G., & McDermott, J. H. (2015). Distinct Cortical Pathways for Music and Speech Revealed by Hypothesis-Free Voxel Decomposition. In Neuron (Vol. 88, Issue 6, pp. 1281–1296). https://doi.org/10.1016/j.neuron.2015.11.035

Obleser, J., Boecker, H., Drzezga, A., Haslinger, B., Hennenlotter, A., Roettinger, M., Eulitz, C., & Rauschecker, J. P. (2006). Vowel sound extraction in anterior superior temporal cortex. Human Brain Mapping, 27(7), 562–571.

Oganian, Y., & Chang, E. F. (2019). A speech envelope landmark for syllable encoding in human superior temporal gyrus. Science Advances, 5(11), eaay6279.

Ohl, F. W., & Scheich, H. (1997). Orderly cortical representation of vowels based on formant interaction. Proceedings of the National Academy of Sciences of the United States of America, 94(17), 9440–9444.

Pelli, D. G. (1997). The VideoToolbox software for visual psychophysics: transforming numbers into movies. Spatial Vision, 10(4), 437–442.

Peterson, G. E. (1961). Parameters of vowel quality. Journal of Speech and Hearing Research, 4, 10–29.

Pineda, L. A., Castellanos, H., Cuétara, J., Galescu, L., Juárez, J., Llisterri, J., Pérez, P., & Villaseñor, L. (2010). The Corpus DIMEx100: transcription and evaluation. Language Resources and Evaluation, 44(4), 347–370.

Pineda, L. A., Pineda, L. V., Cuétara, J., Castellanos, H., & López, I. (2004). DIMEx100: A New Phonetic and Speech Corpus for Mexican Spanish. Advances in Artificial Intelligence - IBERAMIA 2004, 974–983.

Ray, S., & Maunsell, J. H. R. (2011). Different origins of gamma rhythm and high-gamma activity in macaque visual cortex. PLoS Biology, 9(4), e1000610.

Shestakova, A., Brattico, E., Soloviev, A., Klucharev, V., & Huotilainen, M. (2004). Orderly cortical representation of vowel categories presented by multiple exemplars. Brain Research. Cognitive Brain Research, 21(3), 342–350.

Sjerps, M. J., Fox, N. P., Johnson, K., & Chang, E. F. (2019). Speaker-normalized sound representations in the human auditory cortex. Nature Communications, 10(1), 2465.

Strange, W. (1989). Evolving theories of vowel perception. The Journal of the Acoustical Society of America, 85(5), 2081–2087.

Syrdal, A. K., & Gopal, H. S. (1986). A perceptual model of vowel recognition based on the auditory representation of American English vowels. In The Journal of the Acoustical Society of America (Vol. 79, Issue 4, pp. 1086–1100). https://doi.org/10.1121/1.393381

Tang, C., Hamilton, L. S., & Chang, E. F. (2017). Intonational speech prosody encoding in the human auditory cortex. Science, 357(6353), 797–801.

Theunissen, F. E., David, S. V., Singh, N. C., Hsu, A., Vinje, W. E., & Gallant, J. L. (2001). Estimating spatio-temporal receptive fields of auditory and visual neurons from their responses to natural stimuli. Network, 12(3), 289–316.

Yi, H. G., Leonard, M. K., & Chang, E. F. (2019). The Encoding of Speech Sounds in the Superior Temporal Gyrus. Neuron, 102(6), 1096–1110.

